# Targeting cancer-associated cell surface RNAs with oligonucleotide-drug conjugates enables broad antitumor activity

**DOI:** 10.64898/2026.05.08.723688

**Authors:** Cheng Luo, Chen Cheng, Xiaomei Zhao, Jian Zhou, Zhiwen Jing, Qingchuan Ma, Lingyuan Ye, Keyao Zhu, Linfeng Zhao, Han Xu, Hu Zeng

**Author notes:** These authors contributed equally: Cheng Luo, Chen Cheng, Xiaomei Zhao, Jian Zhou.

## Abstract

Recent studies have revealed the presence of RNAs on mammalian cell surfaces, yet the linkage of cell surface RNAs (csRNAs) to cellular states and their potential as extracellular accessible drug targets remains unexplored. Here we develop Cell-surface and Intracellular RNAs Co-mapping (CIRCmap), a highly multiplexed in situ profiling approach that simultaneously detects thousands of cell-surface and intracellular RNAs at single-cell resolution. Using CIRCmap, we uncover that csRNA distributions are correlated with cellular states and identified cancer-associated csRNAs. Integrative analysis of cell-surface and intracellular RNAs within the same cells implies that the csRNAs undergo endocytosis and endolysosomal trafficking, which is further supported by perturbation experiments and co-localization visualization. We design oligonucleotide-drug conjugates (ODCs) that target cancer-associated csRNAs for endocytic delivery of cytotoxic payloads selectively to cancer cells. ODCs exhibit broad antitumor activity in cell lines and an *in vivo* mouse tumor model, opening a promising new avenue for targeted cancer therapy.

## Introduction

Extracellular accessible cell surface components offer distinct advantages as drug targets, with the majority of clinically used drugs targeting membrane proteins^1,2^. A prominent example is the development of antibody-drug conjugates (ADCs), which utilize antibodies to target tumor-associated antigen proteins on the cell surface to deliver highly toxic payloads specifically to cancer cells through endocytosis^3^. ADCs demonstrate enhanced antitumor efficacy compared to conventional chemotherapy and represent one of the fastest-growing anti-cancer drugs in the past decade^4–6^. To date, fifteen ADCs have been approved by the U.S. Food and Drug Administration (FDA), with more than 200 additional candidates in clinical trials^7–9^, establishing targeted endocytic delivery as a transformative modality in oncology.

Recent studies revealed that RNAs are localized on mammalian cell surfaces, where they exhibit biological functions^10–15^. RNAs are an emerging class of druggable targets beyond proteins^16–18^. A key question arises as to whether these cell-surface RNAs (csRNAs) are connected to specific cellular states and if they can be extracellularly accessible drug targets. This is impeded by technical limitations as existing csRNA sequencing methods lack single-cell resolution and cannot simultaneously capture intracellular RNA signals from the same cell^10,11,19^. To address this question, here we developed Cell-surface and Intracellular RNAs Co-mapping (CIRCmap), an integrated approach to simultaneously profile csRNAs and intracellular transcriptomes at single-cell resolution, enabling direct connection between csRNA signatures and cellular states. Using CIRCmap, we demonstrated that csRNA patterns are connected with cellular states and identified specific csRNAs associated with cancer cell types. We further discovered that csRNAs are internalized via endocytosis and are further sorted into late endosomes and lysosomes. Inspired by the similar endocytic uptake mechanism between csRNAs and ADCs, we developed the oligonucleotide-drug conjugates (ODCs) targeting cancer-associated csRNAs for selective cytotoxic payload delivery. We validated the broad antitumor efficacy of ODCs in cell lines with different sources and an *in vivo* mouse tumor model, thereby supporting a novel and promising therapeutic strategy targeting cancer-associated csRNAs.

## Results

### CIRCmap enables joint profiling of cell-surface and intracellular RNAs at single-cell resolution

CIRCmap is built upon a sequential two-round hybridization strategy (Fig. 1a). The first hybridization round occurs prior to permeabilization, while the plasma membrane remains intact, a strategy well used for labeling of cell-surface proteins, glycans and RNAs^11,13,19,20^, ensuring selective targeting of cell surface RNAs by a pair of SNAIL (specific amplification of nucleic acids via intramolecular ligation) probes^21^. Following permeabilization, the second hybridization round enables targeting of intracellular RNAs. The SNAIL probes from both hybridization rounds undergo enzymatic amplification to generate cDNA amplicons, followed by chemical modification and copolymerization within a hydrogel matrix. The RNA-unique barcodes embedded in each DNA amplicon are subsequently decoded via in situ sequencing with error reduction by dynamic annealing and ligation (SEDAL)^21^ (Fig. 1a).

**Fig. 1.**
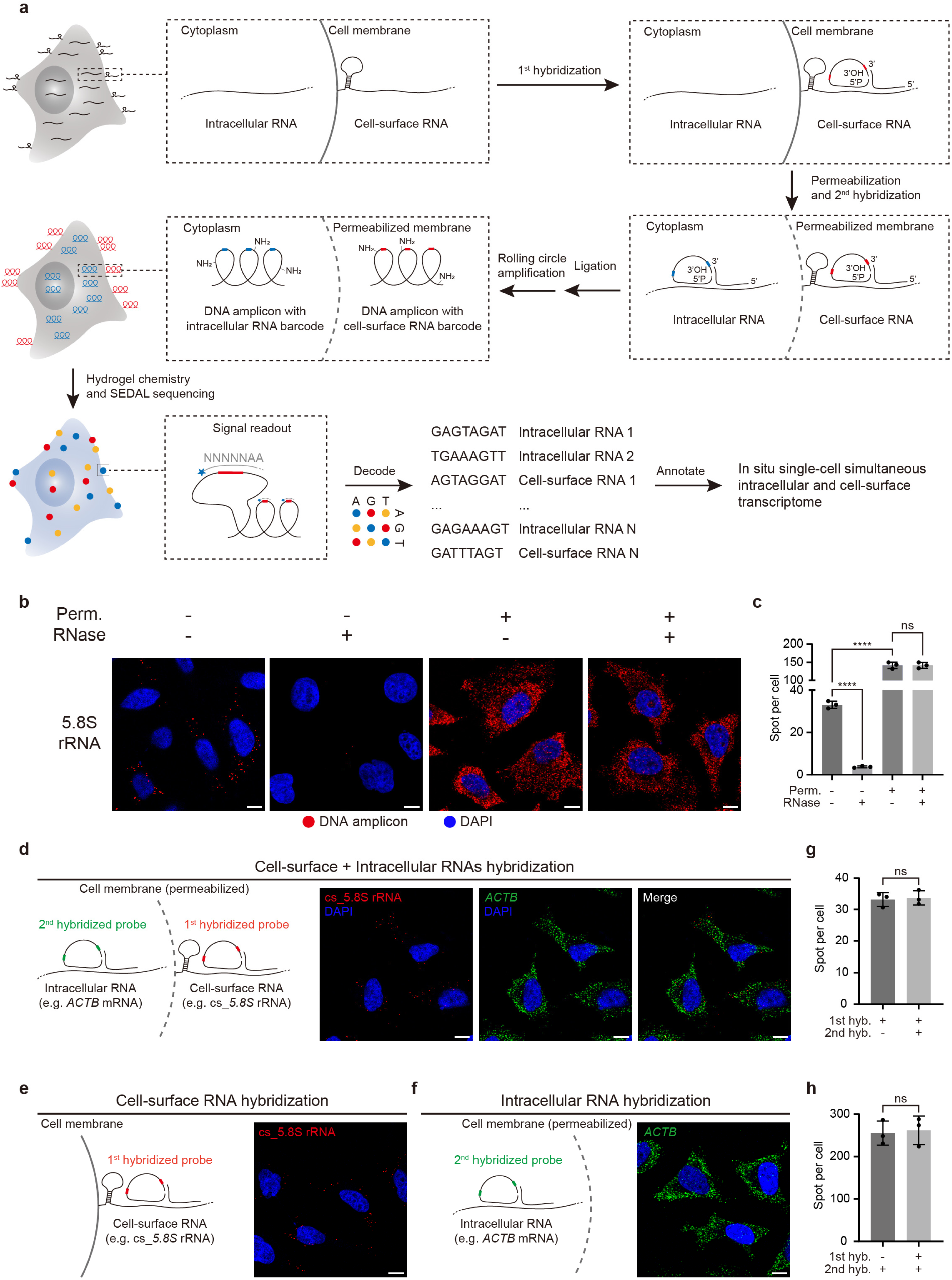
Design and validation of CIRCmap. **a**, Technical schematics of CIRCmap. Followed by the fixation of cells, the 1^st^ hybridization is conducted by hybridizing the paired barcoded padlock and primer probes to csRNAs without cell permeabilization. The 2^nd^ hybridization probes are hybridized to intracellular RNAs after cell permeabilization. Then the ligation reaction results in the intact padlocks serving as the circular templates for rolling circle amplification. Subsequently, the DNA amplicons are incorporated into the hydrogel via copolymerization using tissue-hydrogel chemistry, enabling in situ sequencing. The gene-unique barcode sequence within the DNA amplicons are read out using cyclic in situ sequencing. **b**, Fluorescent images show RNA detection results of 5.8S rRNA under conditions of RNase treatment or not in live cells, permeabilized or not in fixed cells before probe hybridization. The scale bars are 10 μm. **c**, Quantification of the 5.8S rRNA signal spots under conditions shown in panel B. Error bars, standard deviation. n = 3 images per condition. Student’s t-test, ****P < 0.0001, ns, not significant. **d-f**, Diagram and Fluorescent images of cell-surface (cs_5.8S rRNA, red) and intracellular RNA (*ACTB* mRNA, green) detection signal obtained by sequential two-rounds hybridization (D, 1^st^ hybridization of cs_5.8S rRNA, 2^nd^ hybridization of *ACTB* mRNA), 1^st^ hybridization only (E, cs_5.8S rRNA) and 2^nd^ hybridization only (F, *ACTB* mRNA), respectively. The scale bars are 10 μm. **g-h**, The barplot shows the quantification results of cs_5.8S rRNA signal spots (**g**) and *ACTB* mRNA signal spots (**h**) under different hybridization conditions shown in panel (**d-f**). Error bars, standard deviation. n = 3 images per condition. Student’s t-test, ns, not significant.

To validate the reliability of CIRCmap for detecting csRNAs, we chose previously reported glycosylated cell-surface RNAs 5.8S rRNA and U3A (we added “cs” prefix to cell-surface RNA names, e.g. cs_5.8S rRNA and cs_U3A) as representative targets (Supplementary Table 1)^11,19,20^. Probe hybridization was conducted under non-permeabilizing conditions to preserve membrane integrity in line with established csRNA detection methods^19,20^, and DNA amplicon signals generated via in situ amplification were interpreted as detection of csRNAs (Fig. 1b and Extended Data Fig. 1a). Consistent with previous reports^19,20^, pre-fixation treatment of live cells with RNase largely reduced these signals (89% reduction for 5.8S rRNA, 91% reduction for U3A) (Fig. 1c and Extended Data Fig. 1b), confirming that detection under non-permeabilized conditions specifically captures extracellular RNAs. In contrast, when hybridization was conducted after permeabilization, intracellular 5.8S rRNA and U3A were detected with much higher signal spots, and live-cell RNase pretreatment showed no significant reduction in signal intensity (Fig. 1b,c and Extended Data Fig. 1a,b), indicating that extracellular RNase treatment did not interfere with subsequent RNA in situ sequencing. To further verify that the non-permeabilized hybridization strategy avoids detecting intracellular RNAs, we targeted two highly abundant cytoplasmic mRNAs, *ACTB* and *GAPDH*, as controls. As expected, amplicon signals for these mRNAs were only detectable under permeabilized conditions (Extended Data Fig. 1c,d). These validation experiments collectively demonstrate the specificity of CIRCmap for sensitive detection of csRNAs. To assess the orthogonality of the two hybridization rounds, we simultaneously detected cs_5.8S rRNA (via the first round) and *ACTB* mRNA (via the second round). Control experiments using only the first or the second hybridization round showed no significant difference in DNA amplicon signals compared to the full two-round procedure (Fig. 1d,h). This confirms that the two hybridization rounds are non-interfering and demonstrates the capability of CIRCmap for reliable co-detection of cell-surface and intracellular RNAs.

### Multiplexed CIRCmap in cancer and non-tumorigenic cell lines

After benchmarking the specificity and performance of CIRCmap, we applied the method in a multiplexed-gene format across ten cell lines, including five cancer cell lines (HeLa, HepG2, MDA-MB-231, A549, and U2OS) and five non-tumorigenic cell lines (immortalized: hTERT-HPNE and MCF-10A, primary: IMR-90, human dermal fibroblast, and human foreskin fibroblast) (Fig. 2a). For csRNA detection, we targeted non-coding RNA categories based on prior evidence of their cell surface localization^11,12^, including small nucleolar RNA (snoRNA), small nuclear RNA (snRNA), Y RNA, signal recognition particle RNA (SRP RNA), and ribosomal RNA (rRNA), encompassing a total of 688 non-coding RNAs (Supplementary Table 1). We selected 1,455 mRNAs (Supplementary Table 1) to represent the intracellular transcriptome, which are composed of cell-cycle markers, transcription factors, RNA binding proteins, and membrane-transport-related genes^22–27^. We conducted ten rounds of in situ sequencing to detect RNA-unique barcodes, followed by an additional imaging round for organelle staining (STAR methods). To assess the sensitivity and reliability of multiplexed CIRCmap, we first compared per-cell read counts between CIRCmap and the spatial transcriptomics method STARmap and observed comparable results^21,28^ (Extended Data Fig. 1e,f). Next, we performed cross-reference analyses and revealed good correspondence between CIRCmap intracellular transcriptome profiles and public RNA-seq data of the five cancer cell lines^29–32^ (Extended Data Fig. 1g), confirming the robustness of CIRCmap for intracellular RNA detection.To evaluate the specificity of CIRCmap in multiplexed csRNA detection, we compared csRNA profiles of HeLa cells identified by CIRCmap with previously reported glycoRNA datasets and csRNA data from the AMOUR technique^11,19^. This analysis revealed significant overlap between CIRCmap-detected csRNAs and both glycoRNAs and AMOUR-identified csRNAs (Extended Data Fig. 1h). Collectively, These benchmarking results validate the sensitivity and reliability of multiplexed CIRCmap for simultaneous profiling of the intracellular transcriptome and csRNAs.

**Fig. 2.**
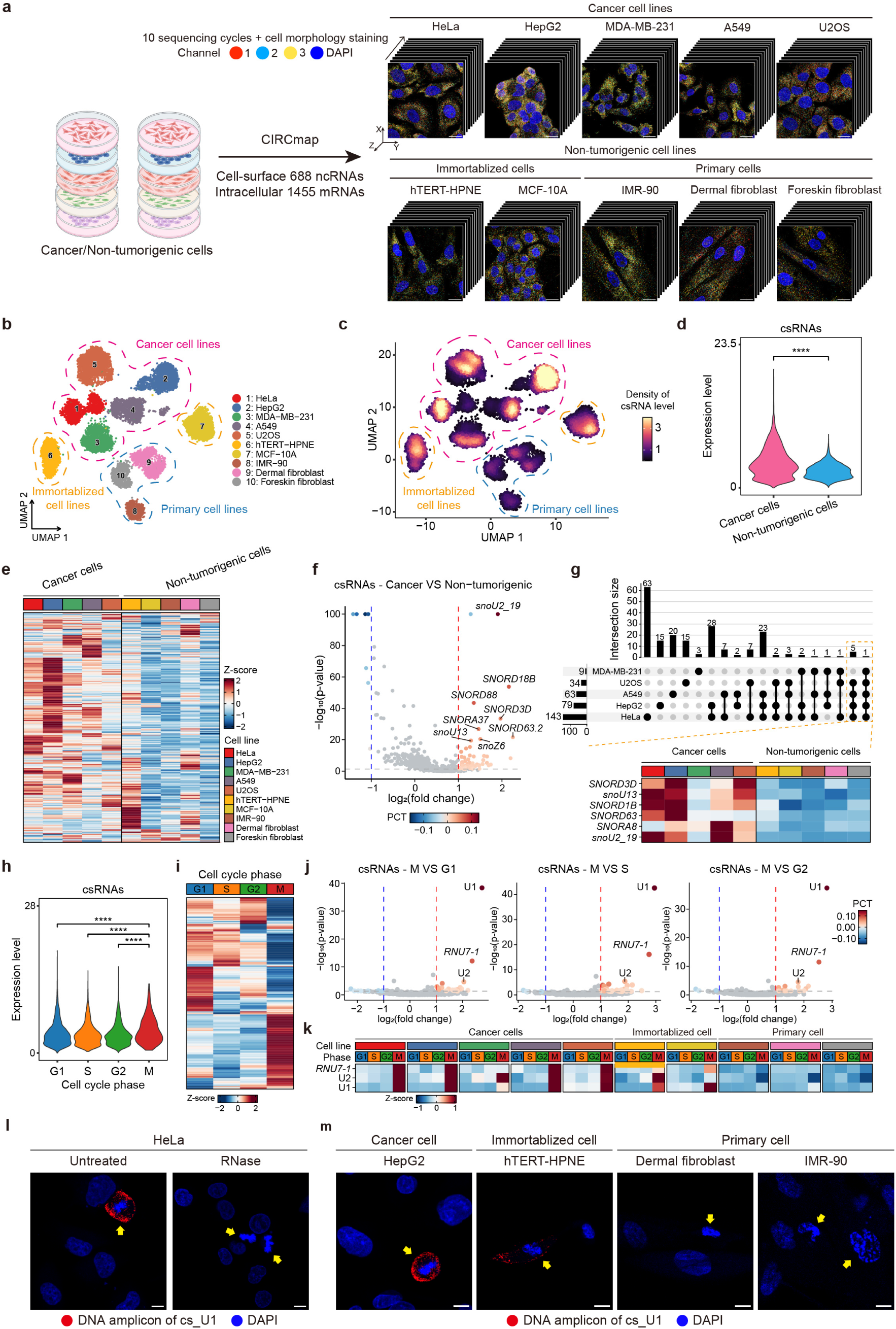
The distribution of csRNAs correlated with cellular states. **a**, Schematic representation of the CIRCmap workflow applied to five cancer cell lines and five non-tumorigenic cell lines. Representative raw images showing the maximum intensity projections of in situ sequencing with zoom-in views of representative cells (right), ten rounds of multiplexed gene unique barcode in situ sequencing to identify 688 csRNAs and 1,455 intracellular mRNAs. **b**, UMAP visualization of ten cell types profiled with mRNAs and csRNAs from CIRCmap result. Cell types are grouped based on their cell source properties (cancerous, immortalized, primary). **c**, csRNA density plot showing csRNA percentage variation across different cell types and cell source properties (cancerous, immortalized, primary). **d**, Violin plot showing the global csRNA expression level in cancer and non-tumorigenic sample groups. Wilcoxon rank-sum test, ns, not significant; ****P < 0.0001. **e**, Heatmap showing the expression pattern of 688 targeted csRNAs across the ten profiled cell lines. **f**, Volcano plots for differential analysis of csRNA levels between cancer and non-tumorigenic sample groups. Differentially detected csRNAs are identified using the Wilcoxon rank-sum test (p value < 0.05 and absolute value of log₂FC > 0.5), and the color of these csRNA dots represents the difference in percentage of cells expressing the csRNA. **g**, UpSet plot showing the intersections and counts of significantly upregulated csRNA sets of the five cancer cell lines compared to the non-tumorigenic sample group (up). Heatmap showing the expression pattern of csRNAs significantly upregulated in at least four cancer cell lines. **h**, Violin plot showing the global csRNA expression level across the four cell cycle phases including G1, G2, S, and M. Wilcoxon rank-sum test, ns, not significant; ****P < 0.0001. **i**, Heatmap showing the expression pattern of 688 targeted csRNAs across the four cell cycle phases. **j**, Volcano plots for differential analysis of csRNA levels between M phase and other three cell cycle phases (G1, G2, S). Differentially detected csRNAs are identified using the Wilcoxon rank-sum test (p value < 0.05 and absolute value of log₂FC > 0.5), and the color of these csRNA dots represents the difference in percentage of cells expressing the csRNA. **k**, Heatmap showing the expression pattern of M phase-enriched csRNAs across the four cell cycle phases in the ten profiled cell lines. **l**, Fluorescent images show cs_U1 detection by single-gene CIRCmap under conditions of RNase treatment or not in HeLa cells. The yellow arrows indicate the M-phase cells. The scale bars are 10 μm. **m**, Fluorescent images show cs_U1 detection by single-gene CIRCmap across multiple cell types, including HepG2 (cancer), hTERT-HPNE (immortalized), dermal fibroblast and IMR-90 (primary) cells. The yellow arrows indicate the M-phase cells. The scale bars are 10 μm.

### csRNAs with cancer-associated distributions

To dissect the specific csRNA patterns at single-cell resolution, we first performed dimensionality reduction analysis of the ten cell lines using Uniform Manifold Approximation and Projection (UMAP), based on the integrated profiling data of 688 csRNAs and 1,455 intracellular mRNAs from CIRCmap (Fig. 2b). As expected, cells were clustered into distinct groups that aligned with their cell identity labels (cancerous, immortalized, primary), indicating the distinct molecular features of different cell types. Visualization of global csRNA density on the UMAP displayed elevated levels in cancer cell lines compared to non-tumorigenic counterparts (Fig. 2c), which was further validated by direct quantitative comparison of total csRNA abundance between cancerous and non-tumorigenic cell lines (Wilcoxon rank-sum test, ****P < 0.0001; Fig. 2d). Notably, cell-surface rRNAs did not exhibit a similar elevation in cancer cells (Extended Data Fig. 2a), suggesting that the enrichment of csRNAs in cancer cell lines is RNA selective rather than a non-specific global increase. We visualized the abundances of 688 targeted csRNAs across the ten cell lines via a heatmap, which revealed the majority of csRNA targets were elevated in cancer cell lines and displayed distinct profiles between cancerous and non-tumorigenic groups (Fig. 2e and Supplementary Table 2). To systematically identify differentially expressed csRNAs, we performed comparative analyses and identified csRNAs with significantly altered levels between cancer and non-tumorigenic cells (Fig. 2f and Supplementary Table 3). For the five cancer cell lines, we individually identified the upregulated csRNA sets compared to non-tumorigenic cells (Extended Data Fig. 2b and Supplementary Table 3), and subsequently interrogated the overlap and uniqueness of these csRNA sets (Fig. 2g). This analysis revealed multiple cell-surface snoRNAs, including *snoU2_19* (ENSG00000199894), *SNORD3D* (ENSG00000277947), snoU13 (ENSG00000238724), *SNORD1B* (ENSG00000199961), *SNORD63* (ENSG00000251987), and *SNORA8* (ENSG00000207410), that were consistently upregulated across diverse cancer types (Fig. 2g and Extended Data Fig. 2c). Together, these results demonstrate a correlation between csRNA distribution and cell types, and reveal csRNAs that are specifically enriched in cancer cells.

We next extended our analysis to explore the correlation between csRNA signatures and cell cycle progression. Prior to csRNA profiling, we established a dual-validation strategy for cell cycle classification: first, intracellular cell cycle marker genes were used to preliminarily assign cells to G1, S, or G2/M phases; second, M-phase cells were further distinguished from the G2/M population based on nuclear morphological features (characteristic chromatin condensation patterns) observed via DAPI staining (Extended Data Fig. 2d). After cell cycle assignment (Extended Data Fig. 2e,f), we then performed quantitative analysis of csRNA levels across cell cycle phases and revealed elevated csRNA levels in M phase compared to G1, S, and G2, respectively (Fig. 2h and Supplementary Table 3), with a notable enrichment of snRNAs (Extended Data Fig. 2g). We next analyzed the expression pattern of all 688 targeted csRNAs across cell cycle phases, uncovering phase-unique csRNA patterns and further showed that M phase exhibited the most prominent csRNA enrichment (Fig. 2i). To identify M-phase enriched csRNAs, we compared csRNA abundances between M phase and each of the other cell cycle phases (G1, S, and G2). This differential analysis revealed a distinct subset of csRNAs specifically associated with M phase, including U1, U2, and *RNU7-1* (Fig. 2j,k). Notably, this M phase enrichment was driven primarily by cancerous and immortalized cells, but not primary cells (Fig. 2k and Extended Data Fig. 2h). Validation via single-gene CIRCmap targeting cs_U1 in HeLa cells confirmed strong M-phase enriched signals, which were abolished upon live-cell RNase treatment (Fig. 2l). Further CIRCmap analysis across different cell types revealed consistent M-phase enrichment of cs_U1 in cancer cell lines (HepG2, A549) and an immortalized cell line (hTERT-HPNE), but not in primary cells (dermal fibroblasts and IMR-90) (Fig. 2M and Extended Data Fig. 2i). To assess the universality of this phenomenon, we performed cs_U1 CIRCmap in other unprofiled cancer cell lines, including human neuroblastoma SH-SY5Y and tumorigenic mouse macrophage RAW 264.7, and observed that both cell lines exhibited M-phase enrichment of cs_U1 signals (Extended Data Fig. 2i).This suggests that M phase-specific csRNA localizations is a widespread feature across cancerous cells from diverse origins and species. These results collectively reveal a correlation between csRNA distribution and cell cycle phases, and identify csRNAs specifically enriched in M-phase of cancerous and immortalized cells.

### csRNAs are endocytosed and undergo endolysosomal trafficking

Leveraging the single-cell joint profiling of csRNAs and the intracellular transcriptome by CIRCmap, we performed correlation analysis to identify genes whose expression levels were associated with csRNA abundances at single-cell level (Extended Data Fig. 3a and Supplementary Table 4). Among the 1,455 intracellular targeted genes, *CAV1* exhibited remarkable negative correlation with global csRNA levels (Fig. 3a). *CAV1* encodes caveolin-1, an essential scaffolding protein that regulates caveolae-mediated endocytosis, facilitating the internalization of specific cargos and membrane components^33^. Along with *CAV1*, *PICALM* (involved in endocytosis)^34,35^ and *RAB5B* (involved in early endosome formation)^33,36^ also exhibited negative correlations with global csRNA levels (Fig. 3a,b). These results suggest that csRNAs are internalized via endocytosis, leading to a reduction in their abundance. To functionally validate this hypothesis, we performed *CAV1* knockdown and overexpression in HeLa cells (Extended Data Fig. 3c), and quantified global csRNA levels via immunostaining using the anti-dsRNA antibody 9D5, as previously described^13^. The csRNA immunostaining signal patterns aligned with the previous study and were susceptible to RNase digestion (Extended Data Fig. 3d,e), confirming the specificity of this csRNA detection approach. We found that *CAV1* knockdown significantly increased the csRNA signal by 34.3% (Fig. 3c,d), while *CAV1* overexpression significantly reduces csRNA signal by 44.8% (Fig. 3e,f), supporting the endocytosis of csRNAs. Interestingly, this finding is consistent with the widely-used ADC target HER2, which exhibits the same tendency upon *CAV1* overexpression and knockdown^37^. In contrast, *RAB35*, a gene that encodes a small GTPase that critically regulates the endosome recycling to control cargo sorting and delivery back to the plasma membrane^38^, showed a positive correlation with csRNA signal. This correlation implies that internalized csRNAs can recycle back to cell membrane (Fig. 3b). To validate this hypothesis, we treated HeLa cells with the endosome recycling inhibitor primaquine, which significantly reduced csRNA signal by 64.2% (Fig. 3g,h), supporting the conclusion that csRNAs undergo endosomal recycling. Notably, cell surface receptor HER2 is similarly affected by primaquine treatment^39^. Together, these results support that endocytosis and endosomal trafficking pathways contribute to csRNA processing, mirroring mechanisms observed for cell surface receptor proteins targeted by ADCs.

**Fig. 3.**
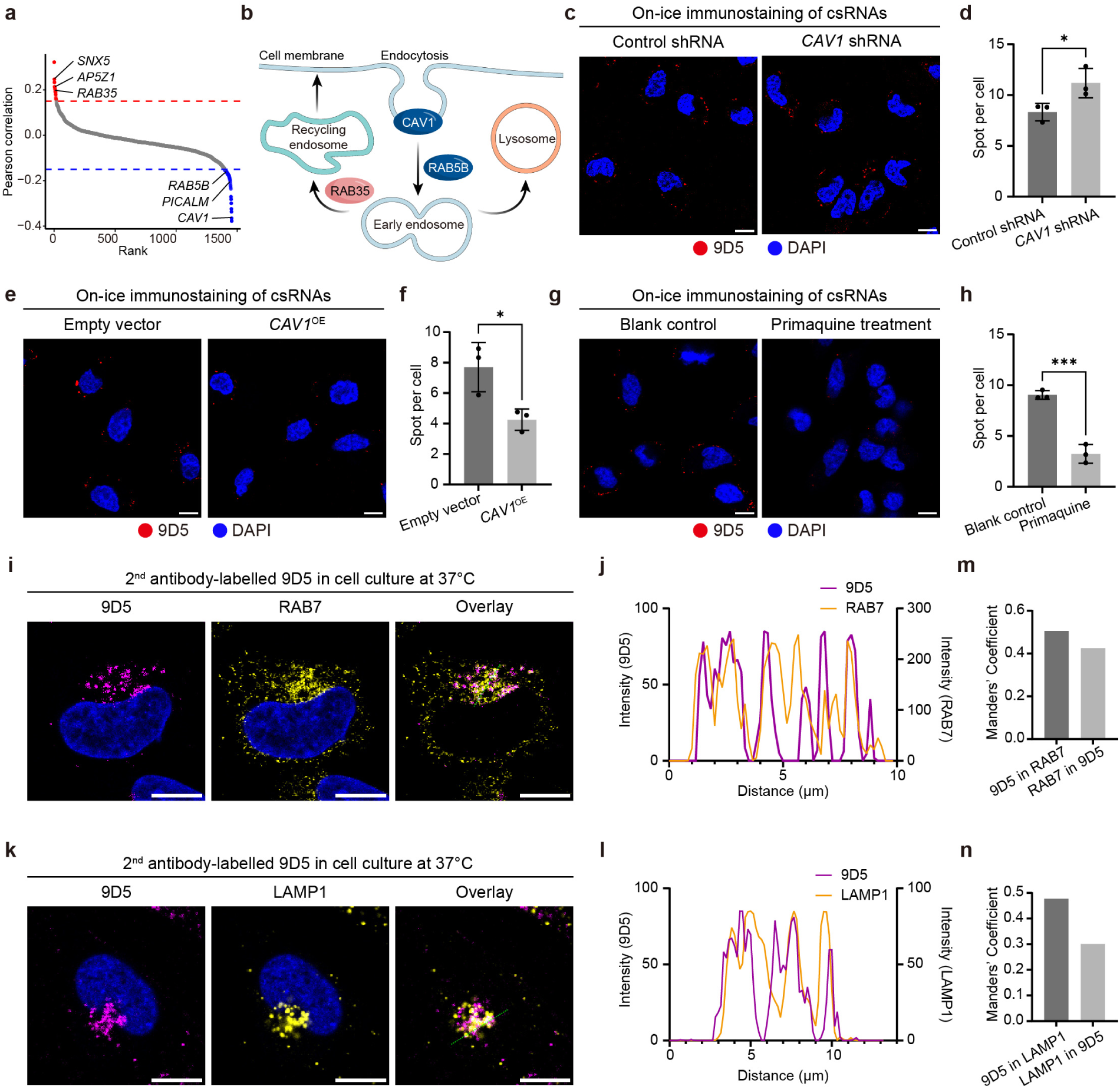
csRNAs undergo endocytosis, endosomal trafficking, and lysosomal entry. **a**, Ranked scatter plot showing the Pearson correlation between the expression levels of the targeted 1,455 mRNAs and global csRNAs in single cells. mRNAs with an absolute value of correlation coefficient > 0.15 are highlighted in red (positive correlation) or blue (negative correlation). **b**, Schematic diagram of endocytosis and endosome trafficking pathways, highlighting genes implicated in these processes and with positive (red) or negative (blue) correlation with global csRNA levels. **c-h**, Fluorescent images and quantitative analysis of csRNA detection results by live-cell immunostaining performed on ice using the dsRNA antibody 9D5 under perturbations, including *CAV1* knockdown (**c**, **d**), *CAV1* overexpression (**e**, **f**) and treatment with endosome recycling inhibitor primaquine (**g**, **h**). Error bars, standard deviation. n = 3 images per condition. Student’s t-test, *P < 0.05, ***P<0.001. The scale bars are 10 μm. **i**, Representative fluorescent images of HeLa cells stained with 9D5 (magenta) in live-cell culture at 37°C, followed by immunostaining of late endosome marker RAB7 (yellow). The scale bars are 10 μm. **j**, The intensity plot of the fluorescence channels across the white line indicated in (**i**). **k**, Representative fluorescent images of HeLa cells stained with 9D5 (magenta) in live-cell culture at 37°C, followed by immunostaining of lysosome marker LAMP1 (yellow). The scale bars are 10 μm. **l,** The intensity plot of the fluorescence channels across the white line indicated in (**k**). **m**, Barplot of Costes-adjusted Manders’ Colocalization Coefficients for the 9D5-RAB7 pair. The fractional overlap of the pair in both directions was calculated and plotted. **n**, Barplot of Costes-adjusted Manders’ Colocalization Coefficients for the 9D5-LAMP1 pair. The fractional overlap of the pair in both directions was calculated and plotted.

To directly visualize the internalized signal of csRNAs, we performed a live-cell assay according to previously described cell internalization assay^13,40^, by culturing cells at 37°C with anti-dsRNA antibody 9D5 labelled with fluorophore-conjugated secondary antibody, then the cell internalization signals are quantified by fluorescence intensity. We observed a 2.7-fold higher 9D5 signal compared to the non-specific internalization signal from an IgG control (Extended Data Fig. 3f,g), revealing the active internalization nature of csRNAs. This 9D5 internalization signal was markedly reduced and exhibited no significant difference from the IgG control upon RNase treatment (Extended Data Fig. 3f,g), further confirming this internalization is csRNA-dependent. Notably, our single-cell mRNA-csRNA correlation analysis also identified highly-correlated genes *SNX5* and *AP5Z1* (Fig. 3a), which are critical for regulating retrograde transport from late endosomes/lysosomes to the trans-Golgi network and maintaining endolysosomal homeostasis^41,42^. These findings further support the involvement of endolysosomal trafficking pathways in csRNA processing. To test this hypothesis, we assessed the colocalization of internalized 9D5 signals with the late endosome marker RAB7 or the lysosome marker LAMP1. Internalized 9D5 signals exhibited clear colocalization with both RAB7 (Fig. 3i,j) and LAMP1 (Fig. 3k,l). To quantify the proportion of internalized csRNAs entering the endolysosomal pathway, we calculated Manders’ coefficients, which demonstrated that 51% and 48% of internalized 9D5 signal overlapped with RAB7 and LAMP1 signals, respectively (Fig. 3m,n). Collectively, these findings provide direct visual evidence that csRNAs undergo endocytosis and endolysosomal trafficking pathway.

### Design of ODCs targeting csRNAs

ADCs represent a breakthrough in targeted cancer therapy by enabling the selective delivery of highly cytotoxic payload to tumors^4–6^. Despite their clinical success, the ADCs face critical limitations including a narrow range of available targets, high manufacturing costs, and the requirement for high-affinity antibodies^4,5,9^. Building on our discovery of cancer-associated csRNAs and their ADC-like endocytic and lysosomal trafficking properties, we developed the ODC platform designed to selectively target csRNAs for drug internalization and release (Fig. 4a). A similar oligonucleotides-based conjugation strategy has been adapted in aptamer-drug conjugates (ApDCs), which use aptamers to target cancer-associated cell surface proteins to deliver therapeutic payloads^43–46^. Pharmacokinetic studies report a half-life (t_1/2_) of approximately ten hours for such oligonucleotide-based conjugates, and *in vivo* efficacy studies validate their antitumor effect and safety across various cancer types^46^, demonstrating the therapeutic potential of this platform. We engineered a proof-of-concept conjugate, termed U1-MMAE, by covalently linking an oligonucleotide probe targeting the well-known csRNA U1 to Monomethyl auristatin E (MMAE), a potent microtubule inhibitor widely used in ADCs, via click chemistry (Fig. 4a). High-resolution capillary electrophoresis confirmed the high conjugation efficiency and purity of the conjugate (Fig. 4b). Immunostaining validation demonstrated successful hybridization of U1-MMAE to csRNAs, as shown by strong colocalization of MMAE signals with csRNA distribution, particularly in M-phase cells where cs_U1 is enriched (Fig. 4c,d). These results confirm the successful synthesis of the ODC and validate its specific targeting to csRNAs.

**Fig. 4.**
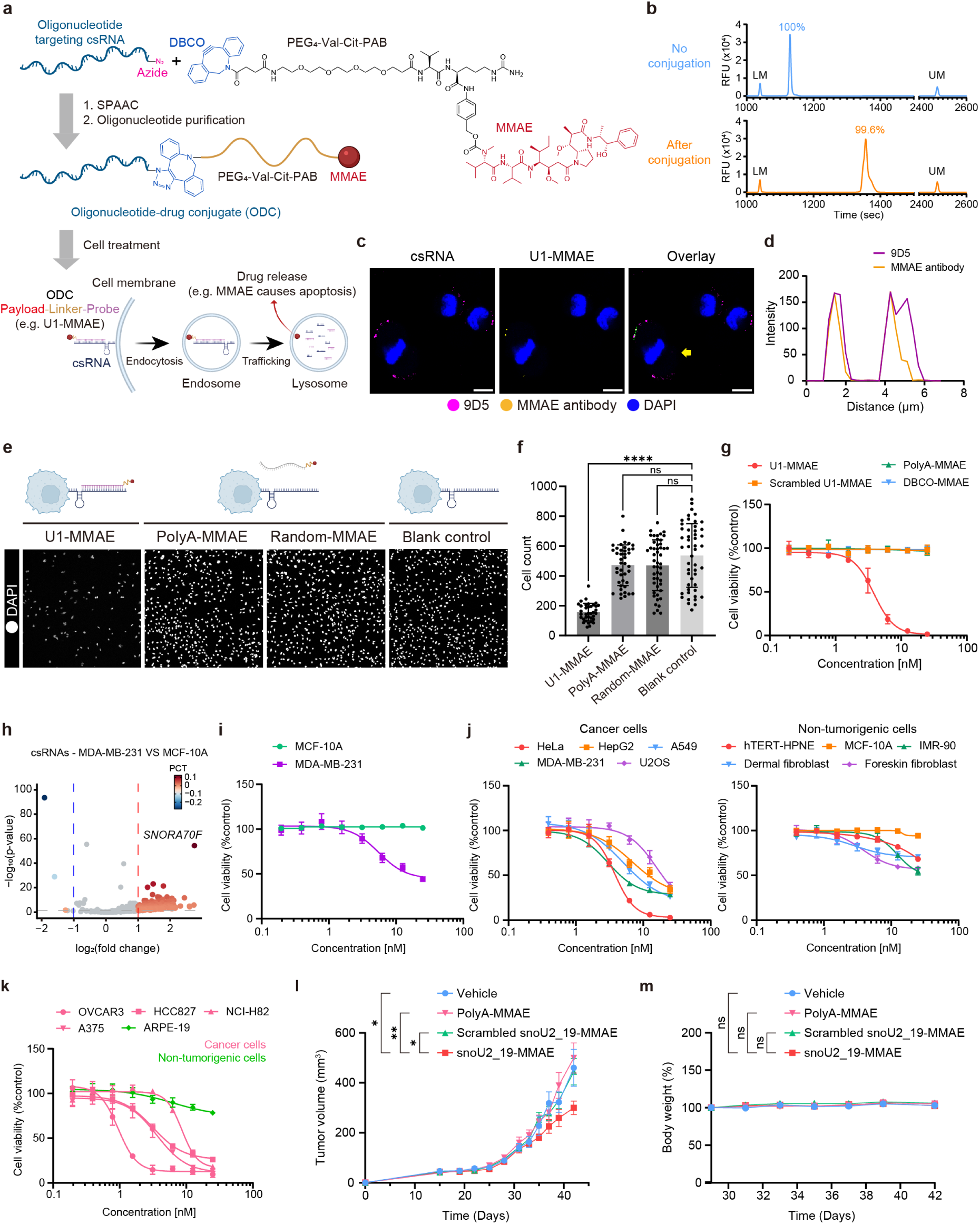
ODCs targeting cancer-associated csRNAs exhibit antitumor activity in cell lines and an *in vivo* xenograft tumor model. **a**, Schematic diagram of ODC manufacturing and its mechanism of action. The ODC is generated by conjugating DBCO-PEG4-VC-PAB-MMAE with 5’ azide modified oligonucleotide via click chemistry. The resulting ODC selectively binds the csRNA target, enabling endocytosis-mediated internalization and subsequent lysosomal release of the drug payload. **b**, Electropherogram (left) and mass spectrum (right) of unconjugated and conjugated U1-targeted oligonucleotides obtained by capillary electrophoresis and LC-MS, respectively. The intensity in the mass spectrum was normalized to the maximum intensity. The standard lower marker (LM) and upper marker (UP) are shown. The purity is defined as the ratio of the product peak area to the total peak area. **c**, Representative immunofluorescence image of HeLa cells treated with U1-MMAE and stained with 9D5 (red) and MMAE antibody (green). The yellow arrows indicate the M-phase cell. The scale bars are 10µm. **d**, The intensity plot of the fluorescence channels across the green dashed line indicated in (**c**). **e**, Representative fluorescent images of HeLa cells under conditions treated with U1-MMAE, polyA-MMAE, random-MMAE or solvent (blank control), followed by fixation and DAPI staining. **f**, Quantification of cell counts per field of view under conditions shown in (**e**). One-way ANOVA, ****P < 0.0001. **g**, Cytotoxicity of U1-MMAE, scrambled U1-MMAE, polyA-MMAE and DBCO-MMAE in HeLa cells after a 72-h incubation. Error bars, standard deviation. n = 3. **h**, Volcano plot for differential analysis of csRNA levels between MDA-MB-231 and MCF-10A cell lines. Differentially detected csRNAs are identified using the Wilcoxon rank-sum test (p value < 0.05 and absolute value of log₂FC > 0.5), and the color of these csRNA dots represents the difference in percentage of cells expressing the csRNA. **i**, Cytotoxicity of SNORA70F-MMAE in MCF-10A and MDA-MB-231 cells after a 72-h incubation. Error bars, standard deviation. n = 3. **j**, Cytotoxicity of snoU2_19-MMAE in the ten CIRCmap-profiled cancer and non-tumorigenic cells after a 72-h incubation. Error bars, standard deviation. n = 3. **k**, Cytotoxicity of snoU2_19-MMAE in additional unprofiled cancer and non-tumorigenic cells after a 72-h incubation. Error bars, standard deviation. n = 2 for ARPE-19, n = 3 for other groups. **l**, Tumor growth curves in HeLa xenograft-bearing mice with intravenous administration of snoU2_19-MMAE, polyA-MMAE, scrambled snoU2_19-MMAE, and vehicle control. Error bars, standard error of the mean. n = 6, unpaired one-tail t-test (day 42), *P < 0.05, **P < 0.01. **m**, Body weight change in HeLa xenograft-bearing mice with intravenous administration of snoU2_19-MMAE, polyA-MMAE, scramble-MMAE, and vehicle control. Error bars, standard error of the mean. n = 6, unpaired one-tail t-test, ns, not significant.

### Antitumor effect of ODCs in cell lines

We then assessed the cytotoxicity and sequence specificity using U1-MMAE in HeLa cells. Treatment with U1-MMAE significantly reduced cell density compared to blank controls (Fig. 4e,f). Crucially, replacing the targeting oligonucleotide with a polyA or random sequence of equal length abolished this cytotoxic effect (Fig. 4e,f), confirming its sequence dependence. To rule out potential artifacts arising from the oligonucleotide or residual DBCO-MMAE, we confirmed that control conjugation products, either lacking DBCO-MMAE in the conjugation mixture or missing the 5′-azide modification on the oligonucleotide, exhibited no detectable cytotoxicity even at markedly higher concentrations (Extended Data Fig. 4a). *In vitro* cytotoxicity assay further demonstrated the high cytotoxic potency of U1-MMAE, yielding an IC₅₀ of 3.83 nM in HeLa cells (Fig. 4g). In contrast, conjugates with non-targeting sequences (polyA or scrambled U1) and free DBCO-MMAE showed minimal reduction in viability (Fig. 4g), reinforcing the sequence-dependent cytotoxicity of U1-MMAE.

To investigate whether ODCs are able to achieve cell-type selective cytotoxicity by targeting specific csRNAs, we first compared csRNA expression between the triple-negative breast cancer cell line MDA-MB-231 and the non-tumorigenic breast epithelial line MCF-10A. Differential analysis identified cs_SNORA70F as the most strongly enriched csRNA in MDA-MB-231 (Fig. 4h). Indeed, the corresponding conjugate SNORA70F-MMAE exhibited significantly greater cytotoxic potency in MDA-MB-231 than in MCF-10A (Fig. 4i). Encouraged by these results, we next engineered snoU2_19-MMAE to target snoU2_19, a csRNA broadly enriched across multiple cancer cell lines (Fig. 2g). To evaluate its potential for broad-spectrum antitumor activity, we treated the five cancer and five non-tumorigenic cell lines (profiled by CIRCmap in this study; Fig. 2a) with snoU2_19-MMAE. The conjugate demonstrated significantly higher cytotoxicity in cancer cell lines compared to non-tumorigenic cell lines (Fig. 4j). Furthermore, we evaluated the potency of snoU2_19-MMAE in additional unprofiled cancer and non-tumorigenic cell lines, where it consistently demonstrated selective toxicity toward cancer cells (Fig. 4k). These results demonstrate the broad antitumor activity of snoU2_19-MMAE. We next investigated whether U1-MMAE could elicit a similarly broad antitumor response, given that cs_U1 is broadly enriched in the M phase of cancer cells but not in primary cells (Fig. 2k). Assessments across both CIRCmap-profiled and additional cell lines confirmed the selective toxicity of U1-MMAE toward cancer cells (Extended Data Fig. 4b,c). Together, these results demonstrate that ODCs targeting cancer-associated csRNAs can achieve broad antitumor activity across diverse cancer cell lines.

### Antitumor effect of ODCs in mouse tumor models

To evaluate the *in vivo* efficacy and safety of ODCs, we first established a xenograft model of human cervical cancer by subcutaneous injection of HeLa cells into immunodeficient mice. Tumor-bearing mice were intravenously administered snoU2_19-MMAE, polyA-MMAE, scrambled snoU2_19-MMAE, or a vehicle control. Treatment with snoU2_19-MMAE led to a significant reduction in tumor volume compared to all other groups (Fig. 4l), demonstrating the sequence-dependent antitumor activity of the ODC *in vivo*. Furthermore, the ODC treatment exhibited no significant toxicity, as evidenced by no significant body-weight loss (Fig. 4m). This safety profile is consistent with the previously reported tolerability of ApDCs, in which MMAE is conjugated to nucleotide aptamers targeting cell-surface proteins^46^.

## Discussion

csRNAs are recently discovered components located on the surface of mammalian cells, offering opportunities as extracellularly accessible drug targets. In this work, we developed CIRCmap for csRNA exploration. To our knowledge, CIRCmap represents the first technology that enabled high-throughput csRNAs profiling at single-cell resolution, as existing high-throughput sequencing approaches, such as Surface-seq and AMOUR, lack single-cell resolution^10,19^, while imaging-based techniques like Surface-FISH, Intact-Surface-FISH, and ARPLA are limited to low gene throughput^10,19,20^. Moreover, CIRCmap simultaneously detected the intracellular transcriptome of the same cells to bridge the gap between csRNA signatures and cellular states. With this technology, we found that cancerous cell lines had higher csRNA abundance than non-tumogenic cell lines and identified csRNAs significantly enriched in cancer cells or M phase of cancer and immortalized cells, highlighting heterogeneity in csRNA distribution across cell types and cell cycle stages and suggesting their functional relevance. The integrative analysis of the csRNA signature and intracellular transcriptome revealed the endolysosomal trafficking of csRNAs, shedding light on the life cycle of csRNAs. Overall, CIRCmap and the accompanying spatial sequencing data from the ten cell types offer a valuable tool and resource for advancing functional studies of csRNAs.

Furthermore, we designed ODCs that target cancer-associated csRNAs, leveraging an endocytic uptake mechanism analogous to ADCs to deliver cytotoxic payloads specifically to cancer cells. We validated the sequence dependent broad antitumor cytotoxicity of these ODCs in both cell lines and mouse tumor models. Compared to ADCs, ODCs utilize short, single-stranded DNA oligonucleotides as targeting moieties and thus offer several key advantages: 1) chemically synthesized DNA oligonucleotides are substantially cheaper and easier to produce than antibodies and allow for straightforward incorporation of chemical modifications^47,48^; 2) the smaller size of ODCs facilitates deeper penetration into tumor tissues, a critical feature for improving therapeutic efficacy in solid tumors^47^; 3) ODCs exhibit lower immunogenic potential^47,48^; minimizing the risk of anti-drug antibody (ADA) responses often associated with protein-based therapies^3^; 4) target recognition via sequence-specific base pairing ensures high selectivity for csRNAs and simplifies the rational design of conjugates, offering a more programmable alternative to the complex protein interactions required for ADC function. Notably, identification of certain csRNA targets, such as cs_snoU2_19, which is broadly enriched across diverse cancer cell lines, and cs_U1, which is consistently elevated during the M phase of cancer cells, indicates their widespread relevance in malignancies. ODCs targeting these csRNAs hold promise for eliciting a broad antitumor response, offering a valuable therapeutic strategy particularly for tumors lacking suitable targets for antibody-based approaches like ADCs. In summary, ODCs exemplify the translational potential of csRNAs as extracellularly accessible drug targets and open new avenues for targeted cancer therapy.

## Methods

### Chemicals and enzymes

Chemicals and enzymes are listed as names (vendor, catalog number): Glass bottom 24-well plates (Cellvis, P24-1.5H-N). 8-well chambered cover glass (Cellvis, C8-1.5H-N). Zymo-Spin V Columns with Reservoir (Zymo Research, C1016-50). 0.45 μm syringe filter (Sigma-Aldrich, SLHPR33RB). Gel slick solution (Lonza, 50640). 3-(Trimethoxysilyl)propyl methacrylate (Sigma-Aldrich, M6514). Poly-D-lysine (Sigma-Aldrich, A-003-M). formaldehyde, 16%, methanol-free (Thermo Fisher Scientific, 28906). Methanol (Sigma-Aldrich, 34860-1L-R). PBS (Thermo Fisher Scientific, 10010-023). Tween-20, 10% (Calbiochem, 655206). Yeast tRNA (Thermo Fisher Scientific, AM7119). SUPERaseꞏIn RNase Inhibitor (Thermo Fisher Scientific, AM2696). Murine RNase inhibitor (Beyotime, R301-03). 20X SSC (Sigma-Aldrich, S6639). Formamide (Calbiochem, 655206). Ribonucleoside vanadyl complex (New England Biolabs, S1402S). T4 DNA ligase, 5 Weiss U/μL (Thermo Fisher Scientific, EL0012). Phi29 DNA polymerase (Thermo Fisher Scientific, EP0094). dNTP mix (New England Biolabs, N0447L). BSA (Sigma-Aldrich, SRE0098-50G). 5-(3-aminoallyl)-dUTP (Thermo Fisher Scientific, AM8439). Methacrylic acid NHS ester, 98% (Sigma-Aldrich, 730300). DMSO, anhydrous (Molecular Probes, D12345). Acrylamide solution, 40% (Bio-Rad, 161-0140). Bis solution, 2% (Bio-Rad, 161-0142). Ammonium persulfate (Sigma-Aldrich, A3678). N,N,N′,N′-Tetramethylethylenediamine (Sigma-Aldrich, T9281). Antarctic phosphatase (New England Biolabs, M0289L). DAPI (Molecular Probes, D1306). Triton X-100, 10% (Sigma-Aldrich, 93443). NaCl (Thermo Fisher Scientific, 91286971). MgCl2 (Thermo Fisher Scientific, AM9530G). Spermidine (Sigma-Aldrich, 85558-1G). Digitonin (Sigma-Aldrich, 300410-250MG). ddNTP (Beyotime, D7401-100μl). Tris-HCl, pH 7.5 (Thermo Fisher Scientific, 15567027). 2-Mercaptoethanol (Sigma-Aldrich, 63689-25ML-F). UltraPure distilled water (Thermo Fisher Scientific, 10977-015). Puromycin (HARVEYBIO, CT0049001). Blasticidin S (HARVEYBIO, PD1513). Polyethylenimine (Polysciences, 23966-100). Polybrene (Beyotime, C0351-1ml). DMEM medium (Thermo Fisher Scientific, C11995500BT). DMEM medium, low glucose, pyruvate (Thermo Fisher Scientific, 11885084). RPMI-1640 medium (Thermo Fisher Scientific, C11875500BT). Ham’s F-12K (Kaighn’s) medium (Thermo Fisher Scientific, 21127022). Fetal bovine serum (Thermo Fisher Scientific, A5669701). MEM non-essential amino acids (Thermo Fisher Scientific, 11140050). Penicillin-Streptomycin (Thermo Fisher Scientific, 15140122). L-Glutamine (Thermo Fisher Scientific, 25030081). RNase A/T1 (Thermo Fisher Scientific, EN0551). RNase I (Thermo Fisher Scientific, EN0601). High-fidelity DNA polymerase (TransGen Biotech, AS231-02). DBCO-PEG4-VC-PAB-MMAE (MedChemExpress, HY-136314). 2,2,2-Tribromoethanol (Sigma-Aldrich, T48402). Small-Scale Total RNA Extraction Kit (Genstone Biotech, TR205). Total RNA Extraction Reagent (ABclonal, RK30129). 2X Universal SYBR Green Fast qPCR Mix (ABclonal, RK21203). 2× MultiF Seamless Assembly Mix (ABclonal, RK21020). Oligo Clean & Concentrator (Zymo Research, D4061). anti-dsRNA (Absolute Antibody, Ab00458-23.0). anti-RAB7 (Santa Cruz Biotechnology, sc-376362). anti-LAMP1 (ABclonal, A21194). anti-MMAE (MedChemExpress, HY-P81056). IgG isotype (Proteintech, 98136-1-RR). anti-mouse IgG Alexa Fluor 488 (Thermo Fisher Scientific, A-21202). anti-mouse IgG Alexa Fluor 546 (Thermo Fisher Scientific, A10036). anti-rabbit IgG Alexa Fluor 546 (Thermo Fisher Scientific, A-10040). anti-rabbit IgG Alexa Fluor 647 (Thermo Fisher Scientific, A-31573).

### Cell culture

HeLa and U2OS cells were obtained from Dr. Qing Li’s laboratory (Peking University) and were cultured in Dulbecco’s Modified Eagle Medium (DMEM) supplemented with 10% fetal bovine serum (FBS) and 1% penicillin-streptomycin (PS). A549, SH-SY5Y and RAW 264.7 cells were obtained from Dr. Yongjun Qian’s laboratory (Peking University). A549 cells were cultured in Ham’s F-12K (Kaighn’s) Medium supplemented with 10% FBS and 1% PS, SH-SY5Y and RAW 264.7 cells were cultured in DMEM supplemented with 10% FBS and 1% PS. IMR-90 cells were obtained from Dr. Jun Liu’s laboratory (Peking University) and were cultured in DMEM (Low Glucose, Pyruvate) supplemented with 10% FBS, 1% PS, 1X L-Glutamine, 1X MEM non-essential amino acids. Human dermal fibroblasts were obtained from Dr. Yang Zhao’s laboratory (Peking University) and were cultured in DMEM supplemented with 10% FBS and 1% PS. Human foreskin fibroblasts were obtained from Dr. Fang Fang’s laboratory (Beijing Children’s Hospital) and were cultured in DMEM supplemented with 10% FBS and 1% PS. HepG2 (HB-8065, ATCC) and GL261 (CVCL_Y003) cells were cultured in DMEM supplemented with 10% FBS and 1% PS. hTERT-HPNE (CRL-4023, ATCC), ARPE-19 (CRL-2302, ATCC), NCI-H82 (HTB-175, ATCC), HCC827 (CRL-2868, ATCC) cells were cultured in RPMI-1640 medium supplemented with 10% FBS and 1% PS. GLMCF-10A (CRL-10317, ATCC) 1:1 DMEM/F12 supplemented with 5% horse serum, 20 ng/mL human epidermal growth factor, 500ng/mL hydrocortisone, 100ng/mL cholera toxin and 1% PS. All cells were cultured at 37°C in a humidified incubator with 5% CO₂.

### Mice

Male C57BL/6 mice (6-8 weeks old) and female NU/J mice (6-8 weeks) were purchased from VitalRiver (Beijing, China). Animals were maintained under specific pathogen-free (SPF) conditions at the Animal Experiment Center, Peking University, Beijing, China. All procedures involving animals followed protocols approved by the Peking University’s Institutional Animal Care and Use Committee (IACUC) and conformed to the Guide for the Care and Use of Laboratory Animals. Besides, the maximal tumor size permitted by the Committee for Animal Research of Peking University is 1500 mm^3^.

### CIRCmap probe design

A systematic pipeline was implemented for the design of CIRCmap padlock and primer probes, proceeding through several sequential stages: (1) Initially, reference transcript sequences were obtained from GENCODE FASTA and GFF3 annotation files, with the Ensembl Canonical transcript selected for protein-coding genes. (2) Each full-length transcript was then segmented into overlapping 40–46 nucleotide candidate regions using OligoMiner^49^, applying specific thresholds for GC content and melting temperature. (3) To minimize off-target binding, these candidate regions were aligned against the reference transcriptome with Bowtie2^50^, retaining only uniquely mapped sequences devoid of variations such as mismatches or INDELs. (4) Repetitive sequences were further filtered by identifying and excluding regions with high-frequency k-mers (k = 18) within the transcriptome using Jellyfish^51^ and OligoMiner. (5) The remaining regions were subdivided into two segments, each 19–25 nucleotides in length, separated by a 0–2 nucleotide spacer. This division was optimized to minimize the melting temperature difference (ΔTm) between segments. (6) To avoid targeting the highly structured regions in non-coding RNAs, the free energies of hybridized regions (sFE) in non-coding RNAs were calculated using RNAfold^52^, and the free energies of probe-RNA hybrids (hFE) were also calculated using NUPACK^53^. The structure scores were recorded as the ratio of hFE to sFE, and this score indicates the ability of probes overcoming the local RNA structure to form hybrids. (7) To address potential secondary structures in probes, the ensemble structure and free energy of each probe were evaluated using the complex analysis module in NUPACK, with probes exhibiting stable secondary structures eliminated. Additionally, the hybridized free energies (hFE) for non-coding RNAs were computed. Finally, a curated set of non-overlapping probes was selected for each gene based on rankings of ΔTm and structure score. This process yielded 1,455 mRNA probes and 688 ncRNA probes, with the majority of RNAs assigned three pairs of primer and padlock probes, and a few of RNAs with limited suitable regions received two or one pair. All probes were synthesized by Synbio Technologies.

### CIRCmap protocol

The cell lines were detached from the culture dishes using TrypLE express enzyme (Gibco) and cultured in the 24-well plates for 1 day before the sample processing. The cells were rinsed with PBS and fixed with 4% methanol-free paraformaldehyde (PFA) in PBS at 37°C for 10 minutes. Then the samples are quenched in 500 μL blocking buffer (0.1 mg/mL yeast tRNA, 0.25 mg/mL BSA, 100 mM glycine, 40 U/mL RNase inhibitor in PBS) at room temperature (RT) for 10 minutes. Next, the 1^st^ hybridization (for cell-surface RNAs) was performed at 37°C for 30 minutes in 200 μL hybridization buffer (50 mM Tris-HCl, 250 mM NaCl, 10 mM MgCl_2_, 0.1 mg/mL yeast tRNA, 0.25 mg/mL BSA, 200 U/mL SUPERaseꞏIn RNase inhibitor, pH 7.5) supplemented with pooled padlock and primer probes (15nM per oligo). Following the hybridization, the samples were washed twice with washing and imaging buffer (10% formamide in 2X SSC) supplemented with 40 U/mL RNase inhibitor, followed by once with PBSR (PBS with 40 U/mL RNase inhibitor), each wash was performed at RT for 10 minutes. The samples were then permeabilized in the permeabilization buffer (0.25 mM spermidine, 0.01% digitonin in PBSR) at 4°C for 20 minutes, RT for 5 minutes. The 2^nd^ hybridization (for intracellular RNAs) was performed at 40°C for 75 minutes in the hybridization buffer supplemented with 10% formamide and pooled padlock and primer probes (15 nM per oligo). Following the hybridization, the samples were washed twice with washing and imaging buffer (10% formamide in 2X SSC) supplemented with 40 U/mL RNase inhibitor, followed by twice with PBSR (PBS with 40 U/mL RNase inhibitor), each wash was performed at RT for 10 minutes. The ligation reaction was performed in the ligation mixture containing 0.05 U/μL T4 DNA ligase, 0.5 mg/mL BSA, and 200 U/mL RNase inhibitor in 1X T4 DNA ligase buffer at room temperature for 2 hours with gentle shaking. Following twice of washes using PBSR at RT for 5 minutes, the rolling circle amplification (RCA) was then conducted by adding 200 μL RCA mixture containing 0.2 U/μL Phi29 DNA polymerase, 250 μM dNTPs, 250 nM ddNTPs, 20 μM 5-(3-aminoallyl)-dUTP, 0.25 mg/mL BSA, and 0.2 U/μL RNase inhibitor in 1X Phi29 buffer, at 30°C for 2 hours. For CIRCmap of a specific gene, the samples can be washed twice with PBS and incubated with a detection mixture (0.2 μM detection probe, 1 μg/mL DAPI, 1X PBS, 1X SSC) at RT for 2 hours, then washed and immersed in PBS for imaging. For multiplexed-gene CIRCmap, the samples were subjected to two PBS washes and then treated with 200 μL of modification solution (25 mM Methylacrylic acid NHS ester in 100 mM sodium bicarbonate buffer) at RT for 1 hour. Next, the samples were briefly rinsed with PBST (0.1% Tween-20 in PBS) and incubated with 150 μL of monomer buffer (containing 4% acrylamide and 0.2% bis-acrylamide in 2X SSC) at room temperature for 15 minutes. After removal of the buffer, 25 μL of polymerization solution (0.2% ammonium persulfate and 0.2% tetramethylethylenediamine in monomer buffer) was carefully applied to the center of each sample, which was swiftly covered with a Gel Slick-coated coverslip. Polymerization was carried out for 1 hour at RT within a nitrogen-filled chamber. Subsequently, the samples were washed twice with PBST for 5 minutes each, followed by incubation in 200 μL of dephosphorylation mixture (0.25 U/μL Antarctic Phosphatase and 0.5 mg/mL BSA in 1X Antarctic Phosphatase buffer) at 37°C for 1 hour. The procedure concluded with three additional PBST washes, each lasting 5 minutes. For SEDAL sequencing, each cycle commenced with three sequential 10-minute incubations in 200 μL of stripping buffer (60% formamide, 0.1% Triton X-100 in water) at RT, followed by three 5-minute PBST washes. Subsequently, samples were incubated for ≥3 hours at RT in 200 μL of sequencing mixture containing 0.09375 U/μL T4 DNA ligase, 0.5 mg/mL BSA, 10 μM reading probe, and 5 μM fluorescent oligos in 1X T4 DNA ligase buffer. Post-incubation, samples underwent three 10-minute washes with 300 μL of washing and imaging buffer (10% formamide in 2X SSC) before imaging in the same buffer.

Imaging was conducted using a Leica STELLARIS 5 confocal microscope equipped with a 40× oil immersion objective (NA 1.3), achieving a voxel resolution of 94.6 × 94.6 × 330 nm. The first ten imaging cycles for in situ sequencing of gene-unique barcodes, enabling detection of 688 cell-surface RNAs and 1,455 intracellular RNAs. After SEDAL sequencing, samples were subjected to three 10-minute stripping buffer treatments and three 5-minute PBST washes. For cell organelle staining, samples were incubated overnight at RT in 200 μL in 1X Flamingo, followed by three PBST washes prior to imaging. DAPI staining was recorded in the first and last round of imaging. For the validation of CIRCmap specificity, RNase-treated samples are prepared by treating live cells with 0.05X RNase A/T1 mix and 0.1 U/μL RNase I for 1 hour before fixation, and permeabilized samples are prepared by applying 500 μL methanol to fixed cells before 1^st^ hybridization followed by incubating at 4°C for 30 minutes, RT for 5 min.

### Image processing and spot finding

The entire analytical workflow was implemented in MATLAB R2023b and Python 3.10.0 and applied according to previous RIBOmap study^28^. First, a modified version of the sparse deconvolution method was implemented to denoise and improve the resolution of spatial imaging^54^. The original Python open-source implementation was adapted to support external parameter input and multi-GPU parallel processing. Sparse deconvolution was performed on HPC cluster equipped with NVIDIA A800 GPUs and a 64-core Intel Xeon processor, using the following parameters: resolution = 250, fidelity = 300, sparsity = 4, deconvolution iteration = 15, chunk overlap = 60, and z-axial continuity = 0.1.

After min-max normalization, images were adjusted by contrast-limited adaptive histogram equalization using the first-round sequencing image as a reference, followed by top-hat filtering to enhance fluorescence signals. Global and local non-rigid registrations were then performed. Global registration used 3D fast Fourier transform to compute cross-correlation across all translational offsets, and the image volume was shifted based on the peak correlation. Non-rigid registration was performed using MATLAB’s *imregdemons()* function, which applies the Demons algorithm to estimate local deformations for precise alignment. After image registration, signal spots were identified in each fluorescence channel of the first sequencing round. Candidate spots were required to meet two criteria: (1) being detected by 3D local maxima extraction (MATLAB *imregionalmax()*, connectivity=26), and (2) exhibiting fluorescence intensities above a predefined threshold (set as 0.2 × the maximum intensity of each channel). For each candidate spot, the maximum fluorescence intensity across the three channels was extracted and used to assign a color for each sequencing round. Spots with ambiguous color identity, defined as having equal maximum intensity in two or more channels, were discarded. The resulting color sequences across rounds were then matched to a predefined gene-specific barcode book to decode gene identities. In CIRCmap design, cell surface RNAs and in-cell RNAs are distinctive on the barcode book. DAPI and Flamingo images were stitched based on the Fourier Shift and cross-correlation optimization. Spatial offsets of each field of view (FOV) were recorded and applied to align the Flamingo channel accordingly. Signal spot stitching was also performed based on the DAPI-derived offsets. Duplicated cells in overlapping tile regions were resolved by retaining the one whose center was closer to the center of the tile.

### Cell segmentation and structured data formatting

For each cell line, deconvoluted Flamingo-stained images were first subjected to maximum intensity projection. The resulting 2D images were then segmented using the Cellpose 3.0.10 via cyto3 pretrained model^55^ with a cell line custom cell diameter to generate 2D reference masks for cell morphology. Similarly, DAPI-stained images after projection were segmented using the Cellpose 3.0.10 via nucleitorch_0 pretrained model to obtain 2D nuclear reference masks.

These segmentation results, together with cell-by-gene count matrices, were represented as AnnData objects using Scanpy (v1.10.2)^56^ and integrated into SpatialData objects (v0.2.1)^57^, enabling unified representation of cell-level expression, transcript-level spatial information, and 2D and 3D cellular and nuclear segmentation data.

### Cell filtering and data preprocessing

In the experiment, residual probe signals were detected in a subset of dead cells. To eliminate the interference from these cells (owing to the disruption of the cell membrane structure, which abolishes the selective permeability of the nuclear envelope to the probes), two key metrics were calculated prior to downstream single-cell analysis: the ratio of total RNA localized within the nucleus, and the proportion of csRNA localized within the nucleus. These metrics were modeled using a multiple linear regression across different cell lines, in which the nuclear proportion of csRNA was regressed on the nuclear-to-total RNA ratio with cell type included as a categorical variable. The linear relationship was fitted using scikit-learn and validated by an F-test with a significance threshold of p < 1 × 10⁻⁵. For cells with abnormally large positive residuals (Z-score > 3) are identified as dead cells due to the non-selective retention of probes caused by compromised nuclear membranes. Count matrices from individual cell lines were integrated into a unified dataset. To ensure data quality, cells with the lowest 5% of read counts were discarded from each cell line. Ultimately, 16,287 cells and 2,143 genes were retained for subsequent analyses.

### Cell cycle identification

Transcriptome-based cell cycle scoring was performed using the *scanpy.tl.score_genes_cell_cycle()* function to classify cells into three cell cycle phases: G1, S, and G2/M^56,58^. Then a custom script was used to associate nuclear regions with individual segmented cell IDs. M phase cells are identified within the transcriptome annotated G2/M phase cells based on their distinct nuclear morphology. Ultimately, all cells were classified into four cell cycle phases: G1, G2, S, and M. The cell cycle labels of the ten cell lines were assigned to the integrated AnnData object.

### Normalization and UMAP visualization

Data normalization was performed using the log1p transformation method integrated in the Scanpy toolkit^56^. Following normalization, 800 highly variable genes (HVGs) were identified to capture biologically meaningful transcriptional heterogeneity. Principal component analysis (PCA) was conducted on the scaled data to reduce dimensionality, after which neighborhood graph construction was performed using *scanpy.pp.neighbors()* with default parameters (n_neighbors=15, metric=’euclidean’), uniform manifold approximation and projection (UMAP) dimensionality reduction was executed via sc.tl.umap(min_dist=1, spread=1) to generate UMAP embeddings for visualization of cellular clustering patterns^59^.

### Differential csRNA analysis

For differential gene expression analysis between distinct cell types and cell cycle phases, we utilized the *FindMarkers()* function via the Wilcoxon rank sum method implemented in the Seurat toolkit (v5.0)^60^. Genes with a log_2_ fold change (logFC) > 1 and p-value < 0.05 were labeled as differentially expressed genes (DEGs), and volcano plots were generated to visualize these DEGs using custom scripts. A csRNA was classified as a “cancer-enriched csRNA” only if it was identified as significantly upregulated (using the thresholds defined above) in at least four out of the five (≥80%) independent cancer cell lines analyzed.

### Cross-reference correspondence analysis

For cancer cell lines in the public datasets, we adopted a unified analytical pipeline: transcriptome quantification was performed with Salmon^61^ against the GENCODE transcriptome reference, followed by normalization to transcripts per million (TPM). Subsequently, a correlation comparison was conducted with cancer cell line data aggregated by cell type using the *AggregateExpression()* function in Seurat.

### csRNA subset enrichment signature calculation

Gene sets corresponding to small nucleolar RNAs (snoRNAs), small nuclear RNAs (snRNAs), Y RNAs, and ribosomal RNAs (rRNAs) were curated based on biotype annotations retrieved from GENCODE (Human Release 47). We applied the AUCell R package to compute csRNA set signature scores, which enabled the comparison of active small RNA expression programs across experimental labels^62^.

### Plasmid construction

For knockdown of *CAV1*, the pLKO.1 plasmids encoding a non-targeting control shRNA (shNC) and shRNAs targeting *CAV1* were obtained from the RNAi Consortium (TRC) collection at the National Center for Protein Sciences (Beijing) - Protein Core, Peking University. The TRC identifiers for the sh*CAV1* constructs are TRCN0000007999 (*CAV1*_1), TRCN0000008000 (*CAV1*_2), and TRCN0000008001 (*CAV1*_3). For overexpression of *CAV1*, the lenti-EF1a-CAV1-3×Flag-P2A-BSD plasmid was constructed by PCR-amplifying the *CAV1* fragment from the template plasmid pCMV-CAV1-3×Flag (RuiBiotech) using a high-fidelity DNA polymerase (TransGen Biotech, AS231-02). The amplicon was subsequently inserted into the linearized lenti-EF1a-MCS-P2A-BSD backbone using the 2× MultiF Seamless Assembly Mix (ABclonal, RK21020).

### Lentiviral preparation and cell transduction

8.0×10^5^ Lenti-X cells were seeded in 6-well plates overnight containing 2 mL DMEM supplemented with 10% FBS. The next morning, 200 μL culture medium was removed and cells were transfected with 0.55 μg pMD2.G (Addgene plasmid #12259), 1.28 μg psPAX2 (Addgene plasmid #12260), and 1.79 μg target plasmid in 200 μL using 9.6 μg PEI. Six hours later, culture medium was completely removed and replaced with fresh DMEM supplemented with 10% FBS. 48 hours after transfection, viral supernatant was harvested and filtered through a 0.45 μm syringe filter (Millipore) and used fresh or kept frozen at -80℃. Cells were seeded 24 hours before transduction to aim for ∼40% confluency at the time of transduction, then were transduced with lentivirus with 8 μg/mL polybrene. For selection of transduced cells, either puromycin or Blasticidin S was added to the culture medium 48 hours post-transduction. Puromycin was used at a final concentration of 1 µg/mL, and Blasticidin S was used at a final concentration of 5 µg/mL. The selection was maintained for 48 hours until all untransduced control cells were dead.

### RNA extraction and qRT-PCR

Total RNA was isolated from cells using Total RNA Extraction Reagent (ABclonal, RK30129) and Small-Scale Total RNA Extraction Kit (Genstone Biotech, TR205). Then, 1 μg of RNA was reverse transcribed using a ABScript III RT Master Mix for qPCR with gDNA Remover (ABclonal, RK20429). qRT-PCR was carried out using 2X Universal SYBR Green Fast qPCR Mix (ABclonal, RK21203) on a Fully Automated Medical PCR Analysis System (TIANLONG, Gentier 96R). Relative gene expression was determined by normalizing the mRNA levels to the reference housekeeping gene *ACTB* using the ddCt method. The primer sequences were as follows: ACTB (forward, 5′-CACCATTGGCAATGAGCGGTTC-3′; reverse, 5′-AGGTCTTTGCGGATGTCCACGT-3′) and CAV1 (forward, 5′-CCAAGGAGATCGACCTGGTCAA-3′; reverse, 5′-GCCGTCAAAACTGTGTGTCCCT-3′). The specificity of the PCR amplification was verified by melting curve analysis of the final products using Medtl System software.

### csRNAs or internalized csRNAs imaging with treatments

The procedure of cell-surface RNA staining with anti-dsRNA antibody (9D5) in live cells is similar to the published work ^13^. Briefly. the WT, KD or OE cells are cultured in 8-well chambered cover glass for 1 day before experiment. For RNase treatment, the cells are treated with 0.05X RNase A/T1 mix and 0.1 U/μL RNase I for 1 hour in cell culture condition; for primaquine treatment, the cells are treated with 100 μM primaquine for 2 hours in cell culture condition. Then the cells are incubated with 5 μg/mL pre-complexed antibodies (9D5:AF647 secondary antibody = 1:2, pre-incubated at 4°C for at least 5h) in blocking solution (0.5% BSA in PBS) on ice for 1 hour. Post-incubation, the cells are washed once with blocking solution, once with PBS, followed by fixation in 3.7% PFA at RT for 15min. Subsequently, the samples are quenched in the blocking solution at RT for 15 minutes. The nucleus staining was done by DAPI staining after the quenching, or by Hoechst 33342 staining before cell fixation. All the samples here are imaged on a Leica STELLARIS 5 confocal microscope.

For the csRNA internalization assay, the HeLa cells are cultured with 40 μg/mL pre-complexed antibodies (9D5/IgG:AF647 secondary antibody = 1:2, pre-incubated at 4°C for at least 5h) in culture medium at 37°C for 12 hours. The RNase treatment was conducted by incubating 0.05X RNase A/T1 mix and 0.1 U/μL RNase I along with the pre-complexed antibodies above. Post-incubation, the cells are washed once with blocking solution, once with PBS, followed by fixation in 3.7% PFA at RT for 15min. Afterward, the samples are quenched in the blocking solution at RT for 1h. The nucleus staining is done by DAPI staining after the quenching, or by Hoechst 33342 staining before cell fixation. For the immunofluorescence of RAB7 and LAMP1, the samples are first incubated with 5 μg/mL primary antibody in blocking solution at 4°C overnight, then at RT for 1 hour, followed by two washes with blocking solution at 4°C for 5 minutes. Next, the 4 μg/mL AF-546 or AF-488 secondary antibodies in blocking solution are applied to samples at 4°C overnight, then at RT for 1 hour. After two washes with blocking solution at 4°C for 5 minutes, the samples are immersed in PBS and ready for imaging.

### ODC synthesis

The 5’ azide oligonucleotides (synthesized by Generay Biotechnology) were prepared at 100 μM in water and stored at -20°C. For the conjugation reaction, SPAAC (strain-promoted azide–alkyne cycloaddition) mixture comprising 20 μM 5’ azide oligonucleotides, 60 μM DBCO-PEG4-VC-PAB-MMAE, 20% DMSO in 1X PBS was prepared and incubated at RT for 8 hours, 37°C for 2 hours. Please note the DBCO-PEG4-VC-PAB-MMAE should be added finally and gently mixed by pipetting to avoid precipitating. Post-incubation, the reacted oligonucleotides were purified by using Oligo Clean & Concentrator (Zymo) following the manual, and stored in 5% DMSO. The purified product can be quantified using NanoDrop One (10-30 μM products were regularly obtained) and qualified using 5200 Fragment Analyzer System and LC-MS (regularly > 95% conjugation ratio). This product can be prepared as aliquots and stored at -20°C until use.

### Cell treatment with ODC

HeLa cells were counted and seeded into 24-well plates at a density of 25,000 cells per well in 500 μL culture medium, followed by a 24-hour incubation to ensure adherence. For ODC treatment, 10 nM ODC was gently mixed with culture medium and added to the wells. After a 72-h culture period, the samples were washed three times with PBS to remove dead cells, fixed with 1.6% PFA at room temperature for 15 minutes, and then stained with 1 μg/mL DAPI for 1 hour. Finally, the samples were washed and immersed in PBS and imaged using a 10× objective lens.

### Cell viability assay

Cells [HeLa, HepG2, MDA-MB-231, A549, U2OS, hTERT-HPNE, MCF-10A, IMR-90, dermal fibroblast, foreskin fibroblast, OVCAR-3, HCC827, NCI-H82, A375, ARPE-19] were plated into culture-treated 96-well clear plates at a density of 5,000 cells per well in 100 μL of culture medium and maintained at 37°C in a humidified incubator with 5% CO₂ for 1 day. After allowing for cell attachment, serially diluted ODCs were administered to each well, and the cells were cultured for an additional 3 days. Upon completion of the treatment period, the spent medium was replaced with 100 μL of WST-8 mixture (10% CCK-8 in culture medium), and the plates were incubated at 37°C for hours according to the manual. Absorbance was then quantified at 460 nm using a BioTek Synergy HTX plate reader. Dose-response curves were fitted, and IC_50_ values were derived through analysis with GraphPad Prism 10 software.

### Mouse xenograft tumor model

Female NU/J mice (6 weeks old) were anesthetized by i.p. injection of 2.5% avertin (2,2,2-tribromoethanol). Hela cells were resuspended in PBS containing 2% FBS and mixed 1:1 (v/v) with Matrigel to obtain a final concentration of 3×10^7^ cells/mL. A total volume of 100 µL of the cell suspension was subcutaneously injected into the right dorsal flank of each mouse using a 1 mL insulin syringe. At day 28 post inoculation, tumor-bearing mice were randomly assigned to four groups (n = 6 per group). snoU2_19-MMAE (1.9 nmol), polyA-MMAE (1.9 nmol), scrambled snoU2_19-MMAE (1.9 nmol) or 5% DMSO in PBS (vehicle control) was administered intravenously at day 29, 31, 33, 35, 37 post tumor inoculation. Tumor volume was monitored 2-3 times per week by measuring the tumor length (a) and width (b) with a caliper, and calculated using the formula V = 0.52×a×b^2^.

## Data availability

All CIRCmap sequenced raw data with the processed version are available on Zenodo (https://doi.org/10.5281/zenodo.17994707). Source data are provided with this paper. All code and analysis are available on GitHub (https://github.com/chengarthur/CIRCmap). All unique materials generated in this study are available from the lead contact with a completed materials transfer agreement.

## Code availability

All code and analysis are available on github https://github.com/chengarthur/CIRCmap.

## Acknowledgements

We sincerely thank Dr. Qing Li (Peking University), Dr. Yongjun Qian (Peking University), Dr. Jun Liu’s (Peking University), Dr. Yang Zhao (Peking University) and Dr. Fang Fang (Beijing Children’s Hospital) for their kind helps in providing cell lines used in this study. We thank the High-Performance Computing Platform of the Center for Life Sciences (Peking University) for the hardware support for data processing. This work is supported by Biomedical Computing Platform of National Biomedical Imaging Center, Peking University. We thank the National Center for Protein Sciences (Beijing)-the Protein Core at Peking University for assistance with the shRNA library and Dr. Tao Xu for technical support. We thank the Peking University High-throughput Sequencing Center (Peking University) for the data acquisition in Agilent 5200 fragment analyzer system. Graphical abstract, Fig. 2a, Fig. 4a, Fig. 4e and Extended Data Fig. 12c include elements created with <u>BioRender.com</u>. This study received support from the National Key R&D Program of China (2024YFF0507402 and 2024YFC3405600 to H.Z.), the National Natural Science Foundation of China (32541046 and 32571669 to H.Z.), the Beijing Natural Science Foundation (Z250014 to H.Z.), Li Ge-Zhao Ning Life Science Youth Research Foundation (LGZNQN202408 to H.Z.), the Frontier Innovation Fund of Peking University Chengdu Academy for Advanced Interdisciplinary Biotechnologies, the Beijing Advanced Center of RNA Biology (BEACON), the Center for Life Sciences (CLS), the College of Future Technology (CFT) of Peking University.

## Author contributions

H.Z. and C.L. designed and developed the CIRCmap. C.L. performed the multiplexed CIRCmap in the ten cell lines. H.X. help to prepare the cell lines. C.C, L.Z. and L.Y. conducted the upstream processing of images. C.C. analyzed the gene expression data, in which H.Z., K.Z., C.L. helped in M-phase cells identification. Z.J. performed *CAV1* knockdown and overexpression in HeLa cells. C.L. conducted the immunostaining experiments in this study. C.L. and H.Z. conceived and developed the ODC. C.L. and Q.M. performed the *in vitro* cytotoxicity assays. X.Z. and J.Z. established the mouse tumor model and conducted the *in vivo* experiments. All authors contributed to writing and revising the manuscript and approved the final version. H.Z. conceptualized and supervised the project.

## Competing interests

H.Z., C.L., X.Z., and Q.M. have filed patent applications related to this work. The other authors declare no competing financial interests. All methods, protocols, and sequences are freely available to nonprofit institutions and investigators.

