## Supplementary Information for "Targeting cancer-associated cell surface RNAs with oligonucleotide-drug conjugates enables broad antitumor activity"

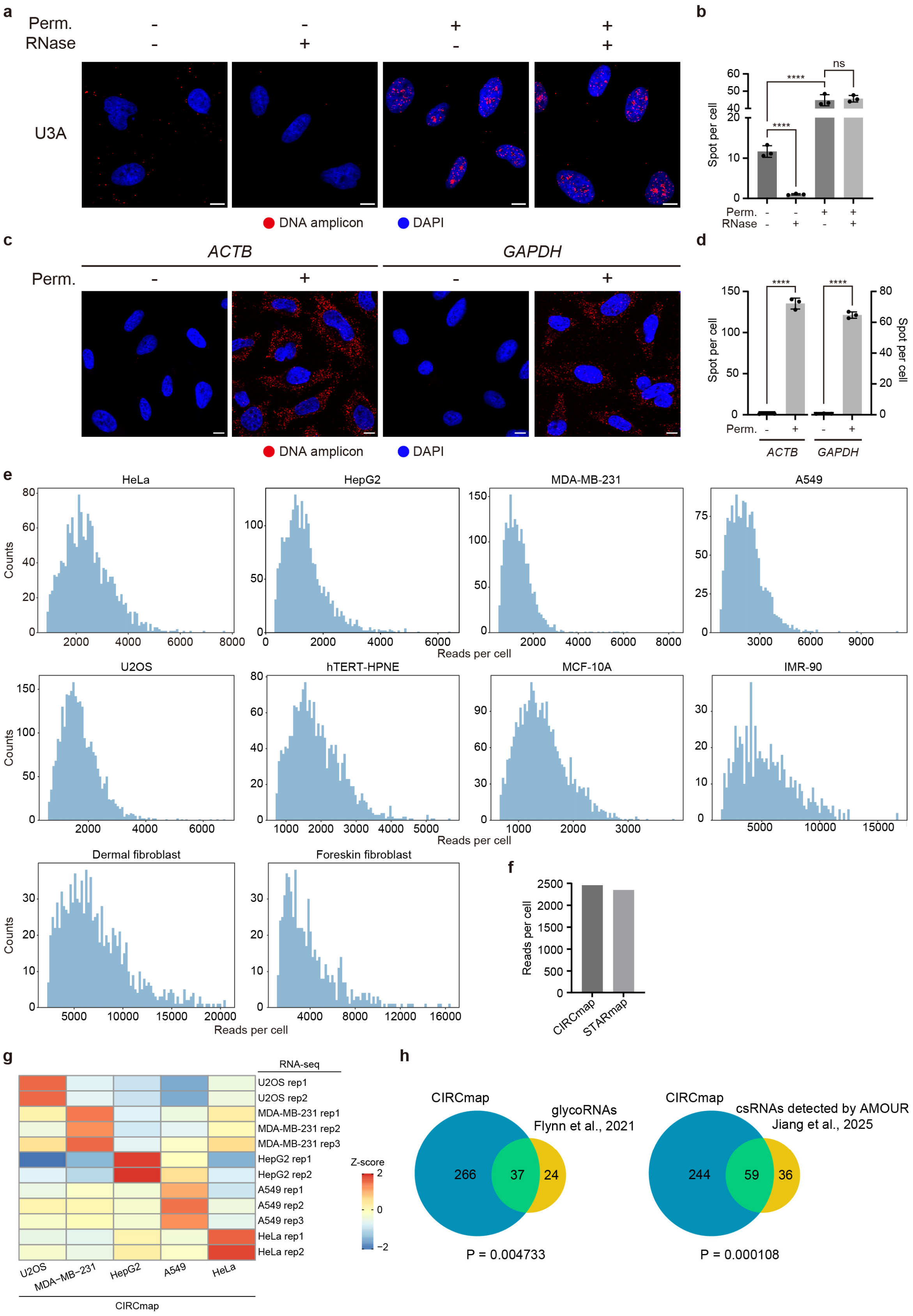
a
Perm.
RNase
U3A
DNA amplicon
DAPI
b
Spot per cell
ns
****
Perm.
RNase
c
ACTB
GAPDH
Perm.
d
Spot per cell
e
HeLa
HepG2
MDA-MB-231
A549
Counts
Reads per cell
U2OS
hTERT-HPNE
MCF-10A
IMR-90
Dermal fibroblast
Foreskin fibroblast
f
Reads per cell
CIRCmap
STARmap
g
RNA-seq
U2OS rep1
U2OS rep2
MDA-MB-231 rep1
MDA-MB-231 rep2
MDA-MB-231 rep3
HepG2 rep1
HepG2 rep2
A549 rep1
A549 rep2
A549 rep3
HeLa rep1
HeLa rep2
Z-score
U2OS
MDA-MB-231
HepG2
A549
HeLa
CIRCmap
h
CIRCmap
glycoRNAs
Flynn et al., 2021
266
37
24
P = 0.004733
CIRCmap
csRNAs detected by AMOUR
Jiang et al., 2025
244
59
36
P = 0.000108

**Extended Data Fig. 1 | Validation and data quality assessment of CIRCmap. a**, Fluorescent images show RNA detection results of U3A RNA under conditions of RNase treatment or not in live cells, permeabilized or not in fixed cells before probe hybridization. The scale bars are 10 μm. **b**, Quantification of the U3A RNA signal spots under conditions shown in (**a**). Error bars, standard deviation. n = 3 images per condition. Student's t-test, ****P < 0.0001, ns, not significant. **c**, Fluorescent images show RNA detection results of ACTB and GAPDH mRNAs in non-permeabilized or permeabilized cells. The scale bars are 10 μm. **d**, Quantification of ACTB and GAPDH signal spots under conditions shown in panel C. Error bars, standard deviation. n = 3 images per condition. Student's t-test, ****P < 0.0001. **e**, Histograms showing the read counts per-cell distribution from CIRCmap across cell lines. **f**, Comparison of average reads per cell between CIRCmap (2,460) and STARmap (2,349) in HeLa cells. **g**, Cross-reference correspondence of CIRCmap intracellular transcriptomes to public RNA-seq datasets of the profiled five cancer cell lines. The color bar represents the normalized Z-score of Pearson correlation coefficient. **h**, Venn diagram showing the overlap of csRNAs detected in HeLa cells using CIRCmap with glycoRNAs reported by Flynn et al. (Cell, 2021) and csRNAs reported by Jiang et al. (Protein & Cell, 2025). The analysis used a background set of all 688 csRNAs targeted by CIRCmap; csRNAs with signals in more than fifty cells are identified as csRNAs detected in HeLa. Statistical significance of overlaps was evaluated via hypergeometric distribution testing.

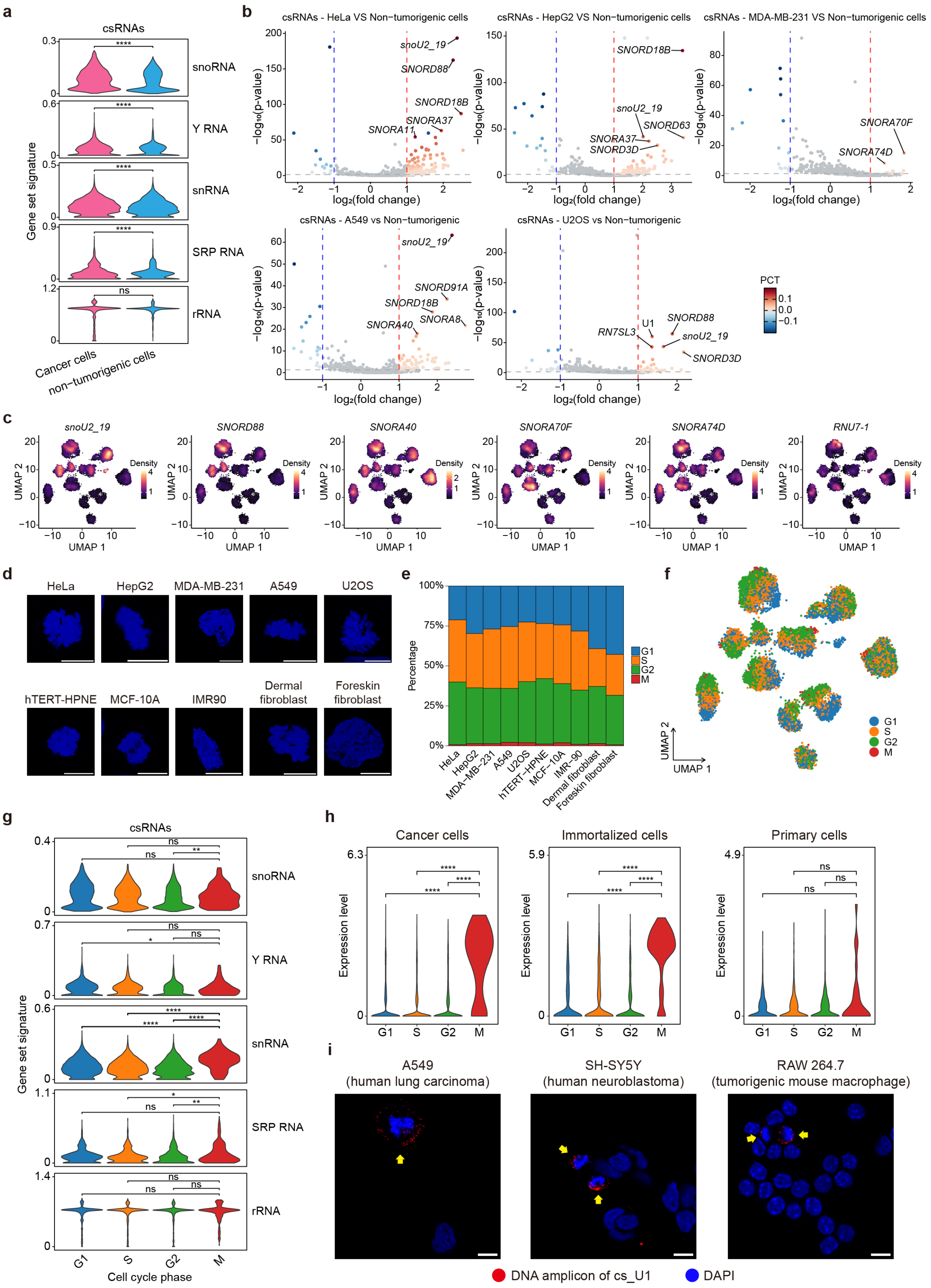
a
csRNAs
snoRNA
Y RNA
snRNA
SRP RNA
rRNA
Gene set signature
Cancer cells
non−tumorigenic cells
b
csRNAs - HeLa VS Non−tumorigenic cells
csRNAs - HepG2 VS Non−tumorigenic cells
csRNAs - MDA-MB-231 VS Non−tumorigenic cells
csRNAs - A549 vs Non−tumorigenic
csRNAs - U2OS vs Non−tumorigenic
−log₁₀(p-value)
log₂(fold change)
snoU2_19
SNORD88
SNORD18B
SNORA37
SNORA11
SNORD63
SNORD3D
SNORA70F
SNORA74D
SNORD91A
SNORA40
SNORA8
RN7SL3
U1
PCT
c
snoU2_19
SNORD88
SNORA40
SNORA70F
SNORA74D
RNU7-1
Density
UMAP 1
UMAP 2
d
HeLa
HepG2
MDA-MB-231
A549
U2OS
hTERT-HPNE
MCF-10A
IMR90
Dermal fibroblast
Foreskin fibroblast
e
Percentage
G1
S
G2
M
HeLa
HepG2
MDA-MB-231
A549
U2OS
hTERT-HPNE
MCF-10A
IMR-90
Dermal fibroblast
Foreskin fibroblast
f
UMAP 1
UMAP 2
G1
S
G2
M
g
csRNAs
snoRNA
Y RNA
snRNA
SRP RNA
rRNA
Gene set signature
Cell cycle phase
h
Cancer cells
Immortalized cells
Primary cells
Expression level
G1
S
G2
M
i
A549
(human lung carcinoma)
SH-SY5Y
(human neuroblastoma)
RAW 264.7
(tumorigenic mouse macrophage)
DNA amplicon of cs_U1
DAPI

**Extended Data Fig. 2 | Distribution patterns of csRNAs across cell types and cell-cycle phases. a**, Violin plot showing csRNA subsets distributions between cancer and non-tumorigenic cell lines. Wilcoxon rank-sum test, ns, not significant; ****P < 0.0001. **b**, Volcano plots for differential analysis of csRNA levels between each individual cancer cell line and non-tumorigenic sample group. Differentially detected csRNAs are identified using the Wilcoxon rank-sum test (p value < 0.05 and absolute value of $\log_2 FC > 0.5$), and the color of these csRNA dots represents the difference in percentage of cells expressing the csRNA. **c**, Density plot showing representative csRNA variation across different cell types. The bars represent the normalized density. **d**, Representative DAPI staining images of M-phase cells from CIRCmap data across all profiled cell lines. The scale bars are 20 µm. **e**, Area plot shows the proportion of different cell cycles (G1, S, G2, M) identified via both transcriptomic and nuclear morphology. **f**, UMAP visualization of the four cell cycle phases from CIRCmap results. **g**, Violin plot showing csRNA subsets distributions across different cell cycle phases (G1, S, G2, M) . Wilcoxon rank-sum test, *P <0.05, **P <0.01, ****P < 0.0001, ns, not significant. **h**, Violin plot showing cs_U1 levels across different cell cycles (G1, S, G2, M), separated by cell source properties (cancerous, immortalized, primary). Wilcoxon rank-sum test, ****P < 0.0001, ns, not significant. **i**, Fluorescent images show cs_U1 detection by single-gene CIRCmap in additional cancer cell types. The yellow arrows indicate the M-phase cells. The scale bars are 10 µm.

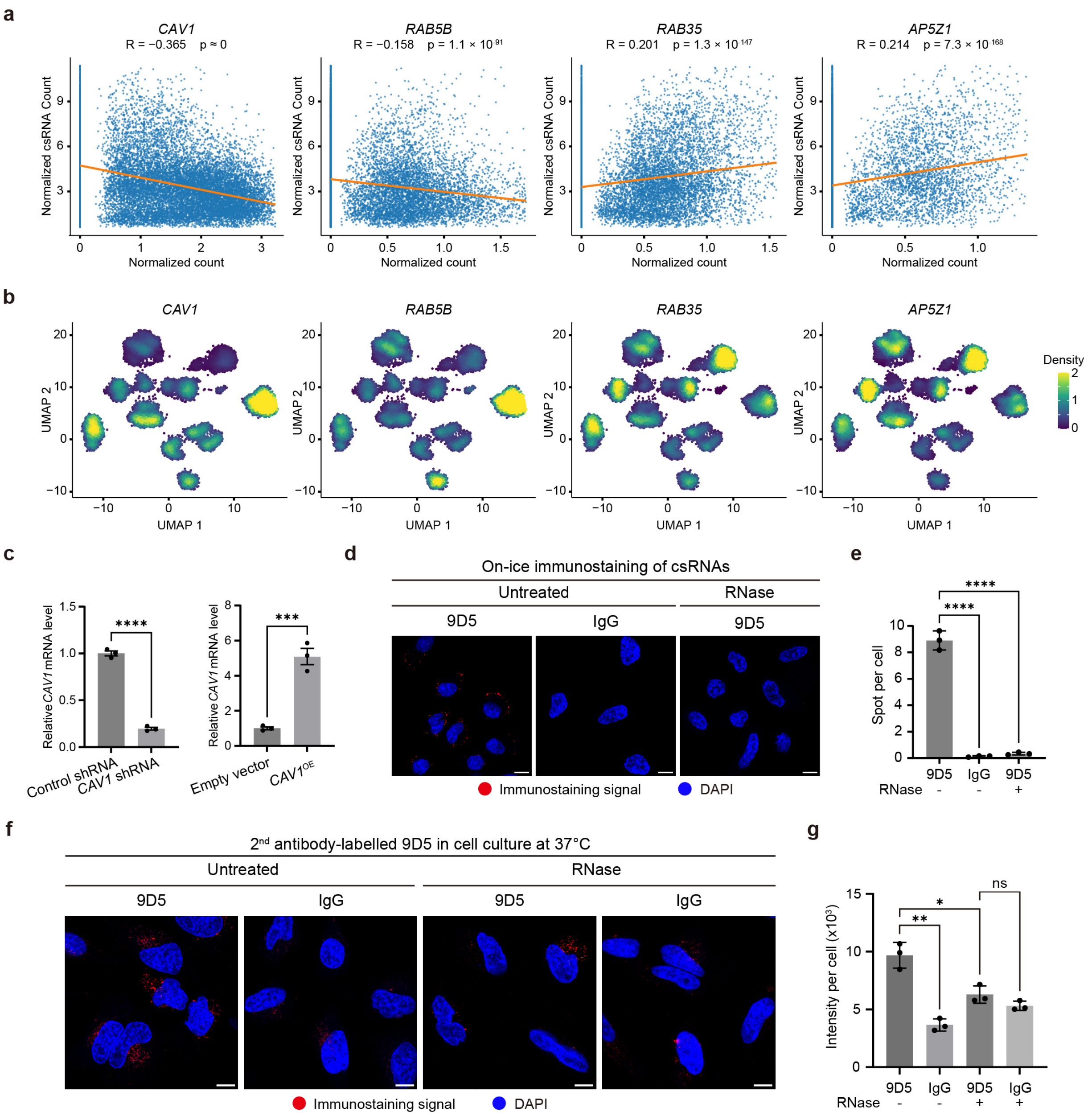


**Extended Data Fig. 3 | Genes associated with csRNA levels and the internalization of csRNAs. a**, Scatter plots showing the correlation between the expression levels of representative mRNAs (*CAV1*, *RAB5B*, *RAB35*, *AP5Z1*) and global csRNA levels (R: Pearson correlation coefficient; p: Pearson correlation test). The orange line indicates the linear regression fit. **b**, Density plot showing variations in representative mRNA expression levels across different cell types. **c**, qPCR quantification of CAV1 mRNA expression levels under overexpression, shRNA knockdown, and control conditions. Error bars, standard deviation. n = 3. Student's t-test, ***$P < 0.001$, ****$P < 0.0001$. **d**, Fluorescence images of live-cell immunostaining performed on ice using the dsRNA antibody 9D5 and IgG control, and 9D5 staining in RNase-pretreated live cells. The scale bars are 10 μm. **e**, Quantification of the immunostaining signal spots under conditions shown in (**d**). Error bars, standard deviation. n = 3. Student's t-test, ****$P < 0.0001$. **f**, Representative fluorescent images of HeLa cells stained with 9D5 or IgG in live-cell culture at 37°C, with or without RNase treatment. The scale bars are 10 μm. **g**, Quantification of the 9D5 and IgG control signal intensity under conditions shown in (**f**). Error bars, standard deviation. n = 3. Student's t-test, *$P < 0.05$, **$P<0.01$, ns, not significant.

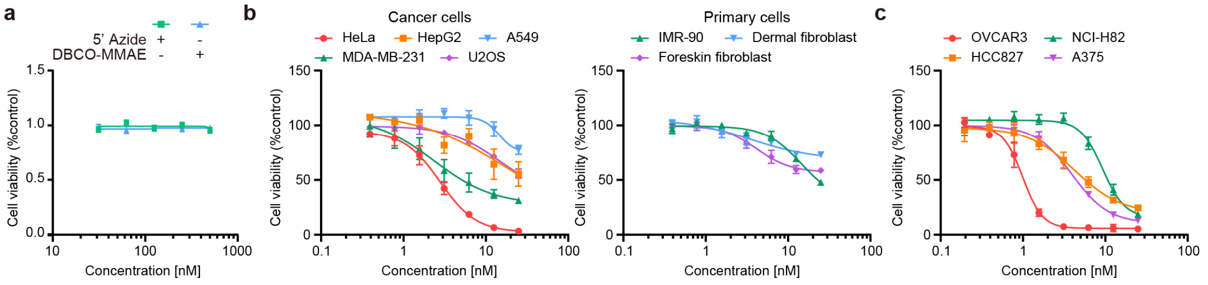


**Extended Data Fig. 4 | Cytotoxicity assay of U1-MMAE in cell lines. a**, Cytotoxicity of U1-MMAE in the CIRCmap-profiled cancer and primary cells after a 72-h incubation. Error bars, standard deviation. n = 3. **b**, Cytotoxicity of U1-MMAE in additional unprofiled cancer cells after a 72-h incubation. Error bars, standard deviation. n = 3. **c**, Cytotoxicity of U1-MMAE in additional unprofiled cancer cells after a 72-h incubation. Error bars, standard deviation. n = 3.
